# *Anopheles* bionomics, insecticide resistance mechanisms and malaria transmission in the Korhogo area, northern Côte d’Ivoire: a pre-intervention study

**DOI:** 10.1101/589556

**Authors:** Barnabas Zogo, Dieudonné Diloma Soma, Bertin N’Cho Tchiekoi, Anthony Somé, Ludovic P. Ahoua Alou, Alphonsine A. Koffi, Florence Fournet, Amal Dahounto, Baba Coulibaly, Souleymane Kandé, Roch Kounbobr Dabiré, Lamine Baba-Moussa, Nicolas Moiroux, Cédric Pennetier

**Affiliations:** Institut Pierre Richet (IPR), Bouaké, Côte d’Ivoire; MIVEGEC, Univ Montpellier, CNRS, IRD, Montpellier, France; Faculté des Sciences et Techniques/Université d’Abomey Calavi, Abomey-Calavi, Benin; Institut de Recherche en Sciences de la Santé (IRSS), Bobo Dioulasso, Burkina Faso; Université Nazi Boni, Bobo-Dioulasso, Burkina Faso

## Abstract

**Background:** A better understanding of malaria transmission at a local scale is essential for developing and implementing effective control strategies. In the frame of a randomized control trial, we aimed to provide an updated description of malaria transmission in the Korhogo area, northern Côte d’Ivoire, and to get baseline data for the trial.

**Methods:** We performed Human Landing Collections in 26 villages in the Korhogo area during the rainy season (September-October 2016, April-May 2017) and the dry season (November-December 2016, February-March 2017). We used Polymerase chain reaction techniques to ascertain the species of the *An. gambiae* complex, *Plasmodium sp* sporozoite infection and insecticide resistance mechanisms in a subset of *Anopheles* vectors.

**Results:** *Anopheles gambiae s.l*. was the predominant malaria vector in the Korhogo area. Overall, more vectors were collected outdoors than indoors (P < 0.001). Of the 774 *An. gambiae s.l*. tested in the laboratory, 89.65% were *An. gambiae s.s*. and 10.35% were *An. coluzzii*. The frequencies of the *kdr* allele were very high in *An. gambiae s.s*. but the *ace-1* allele was found at moderate frequencies. An unprotected individual living in the Korhogo area received an average of 9.04, 0.63, 0.06 and 0.12 infected bites per night in September-October, November-December, February-March, and April-May, respectively.

**Conclusions:** The intensity of malaria transmission is extremely high in the Korhogo area, especially during the rainy season. Malaria control in highly endemic areas such as Korhogo needs to be strengthened with complementary tools in order to reduce the burden of the disease.

## Introduction

Malaria parasites are transmitted to humans by infected female mosquitoes of the genus *Anopheles*, which comprises more than 500 species. About 70 of these species are reported as vectors of human malaria parasites [39]. The intensity of malaria transmission is exceptionally high in Africa largely because of the high vectorial capacity of the major vector species, namely *Anopheles gambiae s.s., Anopheles coluzzii, Anopheles arabiensis* and *Anopheles funestus s.s*. These species display strong anthropophilic host-seeking behavior and have long life expectancy, therefore cause large numbers of secondary malaria cases from one infected individual [42]. The entomological inoculation rate (EIR) that represents the average number of infected bites per person per unit time is generally used to measure the intensity of malaria transmission [40]. The EIR in Africa varies greatly between countries and even within small geographical areas [36]. This heterogeneity needs to be considered when planning and implementing vector control strategies.

Long-lasting insecticidal nets (LLINs) are the main vector control tool used in Côte d’Ivoire to reduce malaria transmission [42]. They are developed to provide physical and chemical barriers against vectors which enter houses to bite humans at night [21]. Consequently, the effectiveness of LLINs may depend on parameters such as the susceptibility of vectors to insecticides and their biting behavior. Unfortunately, resistance to the four insecticide classes used for public health (pyrethroids, organochlorines, carbamates, and organophosphates) is common in the main malaria vector species across Côte d’Ivoire [6,24,37,43]. This resistance involves multiple mechanisms such as metabolic and target site resistance. The *kdr* (responsible for dichlorodiphenyltrichloroethane and pyrethroids resistance) and *ace-1* mutations (responsible for organophosphates and carbamates resistance) are widespread but found at variable frequencies across the country [6,24,37,43]. However, there is very little recent information on the biting behavior of malaria vector species in many areas of Côte d’Ivoire. Studies in several settings across Africa have shown changes in vectors biting behaviors by feeding either mainly outdoors or in the early morning or in the early evening to avoid contact with insecticide-based tools [14,30,33].

Understanding malaria transmission at local scale by determining the main malaria vector species, their abundance, biting behavior, frequencies of insecticide resistance alleles and entomological inoculation rates is a prerequisite for the development and the implementation of effective control and elimination strategies. The present study was conducted as part of a randomized controlled trial (RCT) in the frame of a project called REACT. The project which was conducted in 26 villages in northern Côte d’Ivoire (Korhogo area) and 27 villages in Southwestern Burkina Faso (Diébougou area) was designed to investigate whether the use of vector control strategies in combination with long-lasting insecticidal mosquito nets in areas with intense pyrethroid-resistance provide additional protection against malaria. In this study, we present the results of one year of entomological investigations in the villages of the Korhogo area in Côte d’Ivoire with the aim of providing an updated description of malaria transmission and getting baseline data for the trial.

## Materials and methods

### Study area

The study area included 26 villages of the department of Korhogo (between 9°10’ and 9°40’ N and between 5°20’ and 5°60’ W) located in northern Côte d’Ivoire, West Africa (Figure 1). The villages were selected based on an average population size of 300 inhabitants, a distance between two villages of at least 2 km, and accessibility during the rainy season. The Korhogo area is characterized by a Sudanese climate with a unimodal rainfall regimen from May to November. The annual rainfall varies from 1,200 to 1,400 mm while the mean annual temperature ranges from 21 °C to 35 °C. The minimum temperatures can drop to 16 °C due to the Harmattan wind during December and January. The natural vegetation is mainly a mixture of savannah and open forest characterized by trees and shrubs that are approximately 8-15 m in height. The Korhogo area is fed by tributaries of the Bandama River such as Naramou and Solomougou rivers, which dry up in the dry season. Nevertheless, the area has a high density of water dams that allow agriculture to be practiced throughout the year [17]. Therefore, in areas where the soil is highly conducive to agriculture, most of the local inhabitants are farmers and their staple crops include rice, maize, and cotton. Rice is mainly cultivated during the rainy season in flooded soils although it is also occasionally planted in the dry season in irrigated areas in the vicinity of dams. The main malaria vector control intervention in Korhogo is LLINs which are freely distributed by the government every 3-4 years [34]. The last distribution before this study was performed in 2014.

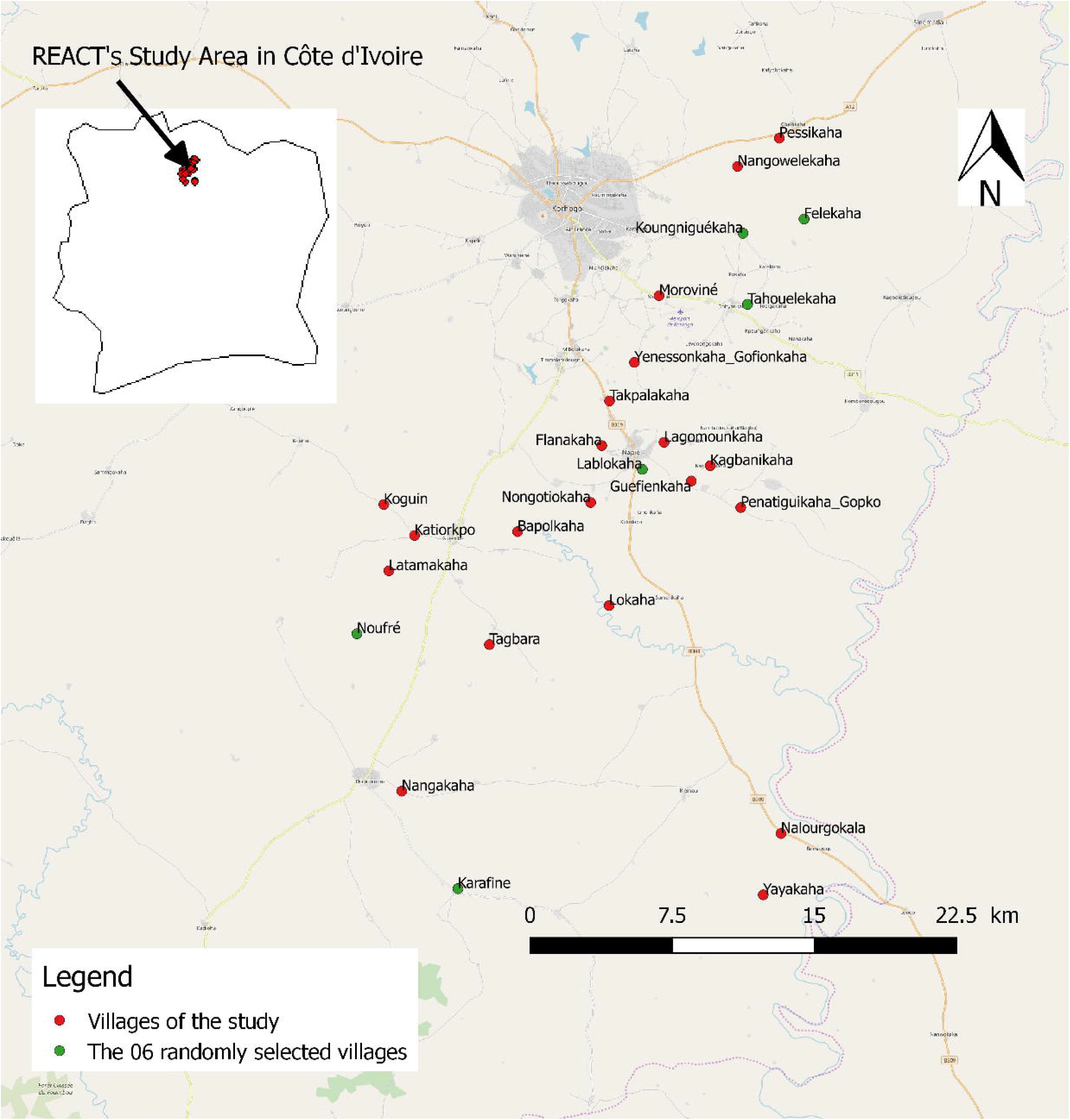

### Mosquito sampling

We performed Human Landing Collections (HLC) in the 26 villages during four surveys: 2 in the rainy season (from September 21^th^ to October 10^th^ 2016 and from April 11^th^ to April 29^th^ 2017) and 2 in the dry season (from November 18^th^ to December 6^th^ 2016 and February 14^th^ to March 04^th^ 2017).

In each village, we collected host-seeking mosquitoes during one night per survey from 17:00 to 09:00 in four randomly selected houses. Each selected house had one collector indoor and one collector outdoor. They were rotated between indoor and outdoor collection sites every hour at each selected house to reduce potential collector bias. Collectors were organized into two teams of 8 persons in each village; the first group collected from 17:00 to 01:00 and the second from 01:00 to 09:00. Each night of collection, one technician from the Institut Pierre Richet (IPR) assisted by two local supervisors supervised the mosquito collections in each village to ensure that they were performing properly.

### *Anopheles* vector processing

After collections, we morphologically identified in the field all the mosquitoes to species or species complex level according to established taxonomic keys [16,17]. Only *Anopheles spp*. mosquitoes were kept and were stored individually in labeled tubes with silica gel at −20°C for molecular analysis. Due to the very large numbers of vectors collected, we analyzed in the laboratory a subsample of *Anopheles spp*. vectors from 6 villages (Tahouelékaha, Koungniguékaha, Lablokaha, Felékaha, Noufré, and Karafiné) randomly chosen out of the 26 villages included in the study (Figure 1). From the vectors collected in these 6 villages, we randomly selected one individual of *Anopheles gambiae s. l*. per hour per collection site during each survey. We analyzed all the *Anopheles nili* complex individuals and all the members of the Funestus Group collected in the 6 villages. We extracted DNA from the head and thorax of each specimen of the subsample using the Livak method [26]. We identified sibling species of the *Anopheles gambiae* complex by Polymerase Chain Reaction (PCR) [13,38]. Quantitative PCR (qPCR) was performed to investigate the presence of the *kdr* L1014F and *ace-1* G119S mutations in the individuals belonging to the *Anopheles gambiae* complex [2,12] and to screen all the selected *Anopheles spp*. mosquitoes for *Plasmodium falciparum* sporozoite infection [4].

### Ethical approval

The ethical clearance for the study was granted from the national ethical committee (N° 063/MSHP/CNER-kp). We received community agreement before the beginning of the study and we obtained written informed consent from all the mosquito collectors and supervisors. Yellow fever vaccines were administered to all the field staff. Collectors were treated free of charge when they were diagnosed with malaria during the study period. They were also free to withdraw from the study at any time without any consequences.

### Data analysis

The human biting rate (HBR) for all malaria vector species was calculated as the number of *Anopheles* vectors (An. *gambiae s.l*., *An. funestus* group, *An. nili* complex) collected per man per night. The *P. falciparum* sporozoite rate in each vector species population was calculated as the number of vectors positive for sporozoites over the number of vectors tested. For each survey, the daily entomological inoculation rate (EIR) was calculated by multiplying the mean human biting rate for all malaria vector species by the *P. falciparum* sporozoite rate in all malaria vector species. Statistical analyses were performed using R software [34].

In order to compare HBRs between surveys and between collection position (indoors or outdoors), we analyzed counts of *Anopheles* vectors (all species) using a negative binomial mixed effect model (function ‘glmer’ from the package lme4) [5]. The fixed variables were the survey, the collection position (indoors/outdoors) and their interaction. Villages and collection houses were used as random intercepts.

We compared the sporozoite rate between surveys, species and collection position (indoor/outdoor) using a binomial mixed effect model (function ‘glmer’ from the package lme4). The fixed variables were the surveys, the *Anopheles* species and the collection position (indoors/outdoors). Villages were set as random intercepts.

For both the HBR and SR models, we performed backward stepwise deletion of the fixed terms followed by Likelihood ratio tests. We used the Tukey’s post-hoc test to compare *Anopheles* counts among surveys according to the final model. Rates Ratio, Odds-Ratio and there 95% confidence interval were computed.

In order to compare night’s biting activity profiles of *An. gambiae s. l*. among surveys, we compare the distribution of collection times using a Kruskall-wallis rank sum test followed by a multiple comparison Dunn’s post-hoc test.

We used the G-test [18] implemented in Genepop 4.7 and run in R [15] to compare gene frequencies at the *kdr* and *ace-1* loci between *An. gambiae* s.s. and *An. coluzzii*, among surveys and among villages [35]. We tested the distribution of genotypes at the *kdr* and *ace-1* loci for conformity to Hardy-Weinberg equilibrium within both species of *An. gambiae* complex, in each survey and in each village, using exact tests as implemented in GenePOP [35].

## Results

We collected a total of 31 324 female mosquitoes belonging to 5 genera (*Anopheles, Culex, Aedes, Mansonia*, and *Uranotania*) and 23 species or species complexes/groups in the 26 villages during the 4 surveys. *Anopheles gambiae s.l*. represented a large proportion of the mosquito species collected, accounting for 94.86%, 59.76%, 74.68% and 92.12% of the total collected in September-October 2016, November-December 2016, February-March 2017 and April-May 2017, respectively. All the mosquitoes identified as members of the *Funestus* Group and the *An. nili* complex represented less than 2% of the total mosquito collected (Table 1). Speciation was successful for 97% of the 764 *An. gambiae s.l*. genotyped to species by PCR. Out of the 356 *An. gambiae s.l*. collected indoor and successfully tested, 314 (88.20%) were *An. gambiae s.s*. and 42 (11.80%) were *An. coluzzii*. Of the 388 *An. gambiae s.l* collected outdoor and successfully tested, 353 (90.98%) were *An. gambiae s.s*. and 35 (9.02%) were *An. coluzzii*. In total, 89.65% of *An. gambiae s.l*. collected and successfully tested were *An. gambiae s.s*. and 10.35% were *An. coluzzii* (Table 2).

**Table 1:**
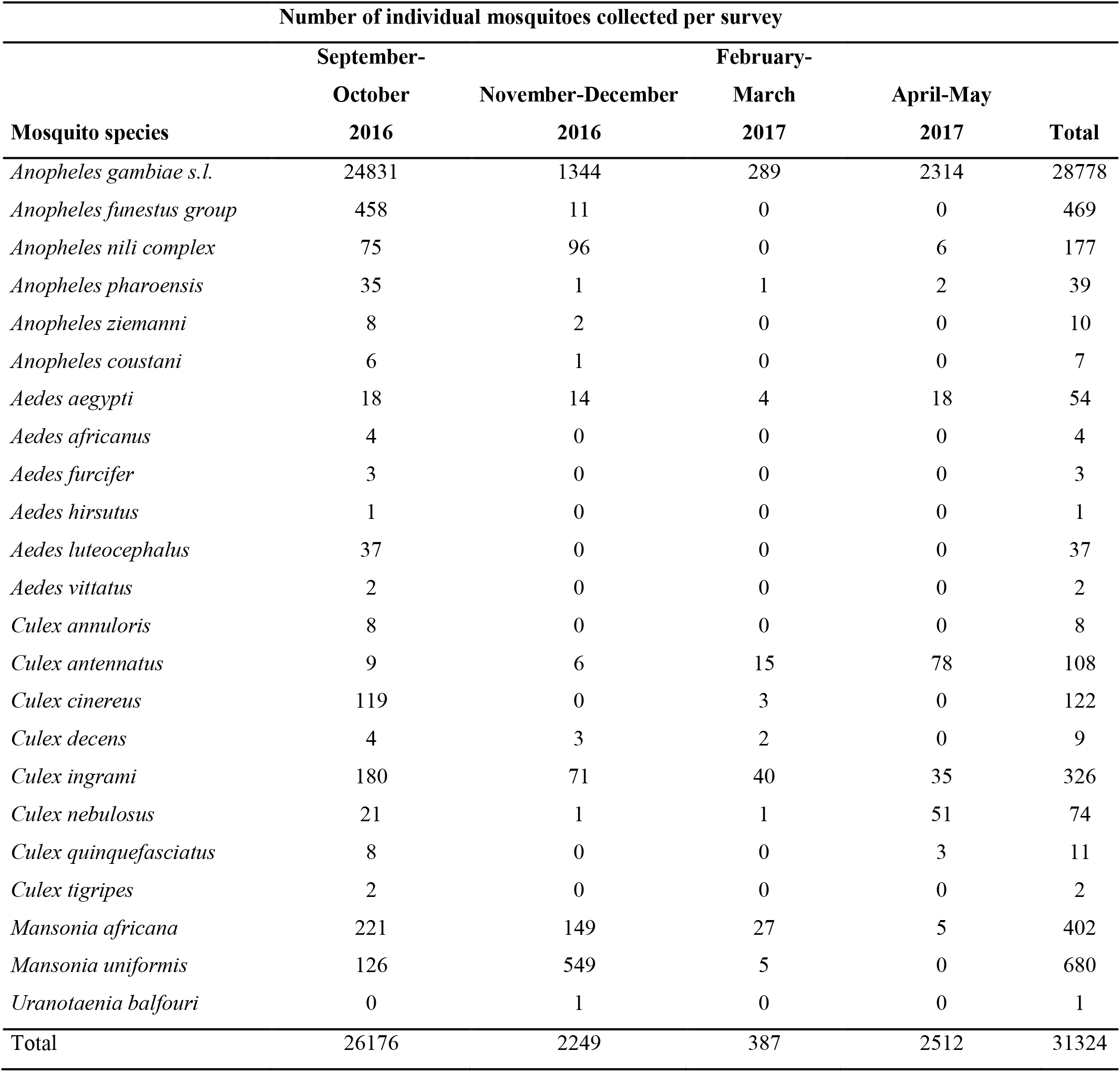
Diversity and abundance of mosquito species in the 26 villages in the Korhogo area during the four surveys

**Table 2:** *Anopheles gambiae* complex species in a subset analyzed

| Survey | Species | No individuals collected |  |  |
| --- | --- | --- | --- | --- |
|  |  | Indoor | Outdoor | Total |
| September-October | *Anopheles gambiae* | 197 | 211 | 408 |
|  | *Anopheles coluzzii* | 8 | 2 | 10 |
|  | **Total** | **205** | **213** | **418** |
| November-December | *Anopheles gambiae* | 74 | 86 | 160 |
|  | *Anopheles coluzzii* | 3 | 2 | 5 |
|  | **Total** | **77** | **88** | **165** |
| February-March | *Anopheles gambiae* | 19 | 24 | 43 |
|  | *Anopheles coluzzii* | 11 | 15 | 26 |
|  | **Total** | **30** | **39** | **69** |
| April-May | *Anopheles gambiae* | 24 | 32 | 56 |
|  | *Anopheles coluzzii* | 20 | 16 | 36 |
|  | **Total** | **44** | **48** | **92** |

The mean HBR for all malaria vector species (An. *gambiae s.l., Funestus* Group, *An. nili* complex) in the study area was 35.37 bites per human per night (interquartile range (IQR): 0.00-42.25) during the four surveys. The mean HBR for all malaria vector species was 121.94 (IQR: 68.50–156.50), 6.98 (IQR: 1.00-8.25), 1.39 (IQR: 0.00-1.00) and 11.15 (IQR: 0.00-9.00) bites per human per night in September-October 2016, November-December 2016, February-March 2017 and April-May 2017, respectively. According to the count model, the HBR for all malaria vector species varied significantly between surveys (χ^2^ = 1746.8, df = 3, P<0.001). It was significantly higher in September-October 2016 than in November-December 2016 (OR [95%CI] = 23.82 [18.78, 30.20]), February-March 2017 (OR = 208.60 [155.25, 280.28]) and April-May 2017 (OR = 28.91 [22.07, 37.87]).

## Biting patterns of malaria vector populations

More vectors were collected outdoors than indoors (OR [95%CI] = 1. 26 [1.16-1.37], P < 0.001) and the outdoor biting rate did not vary significantly between the surveys (χ^2^ = 3.24, df = 3, P=0.3693). During the four surveys, indoors and outdoors biting activities of *An. gambiae s.l*. occurred from dusk to dawn. Both indoors and outdoors biting activities of *An. gambiae s.l*. peaked between 04:00 and 05:00 in September-October 2016 and between 02:00 and 03:00 in November-December 2016 (Figures 2A and 2B). In February-March 2017, the majority of indoor and outdoor bites by *An. gambiae s.l* occurred between 01:00 and 04:00 and between 03:00 and 04:00, respectively (Figure 2C). In April-May 2017 survey, both indoors and outdoors biting activity curves of *An. gambiae s.l*. showed a bimodal shape. Indoors, there was one peak between 01:00 and 02:00 and a smaller one between 04:00 and 05:00. Outdoors, there was one peak between 00:00 and 01:00 and a bigger one between 04:00 and 05:00 (Figure 2D). Biting activity profiles of *An. gambiae s.l*. differed significantly between surveys (Kruskal-Wallis chi-squared = 217.76, df = 3, p-value < 0.001) with activity during the first survey (September-October 2016) peaking later than during the other surveys (Dunn Test p-values < 0.001).

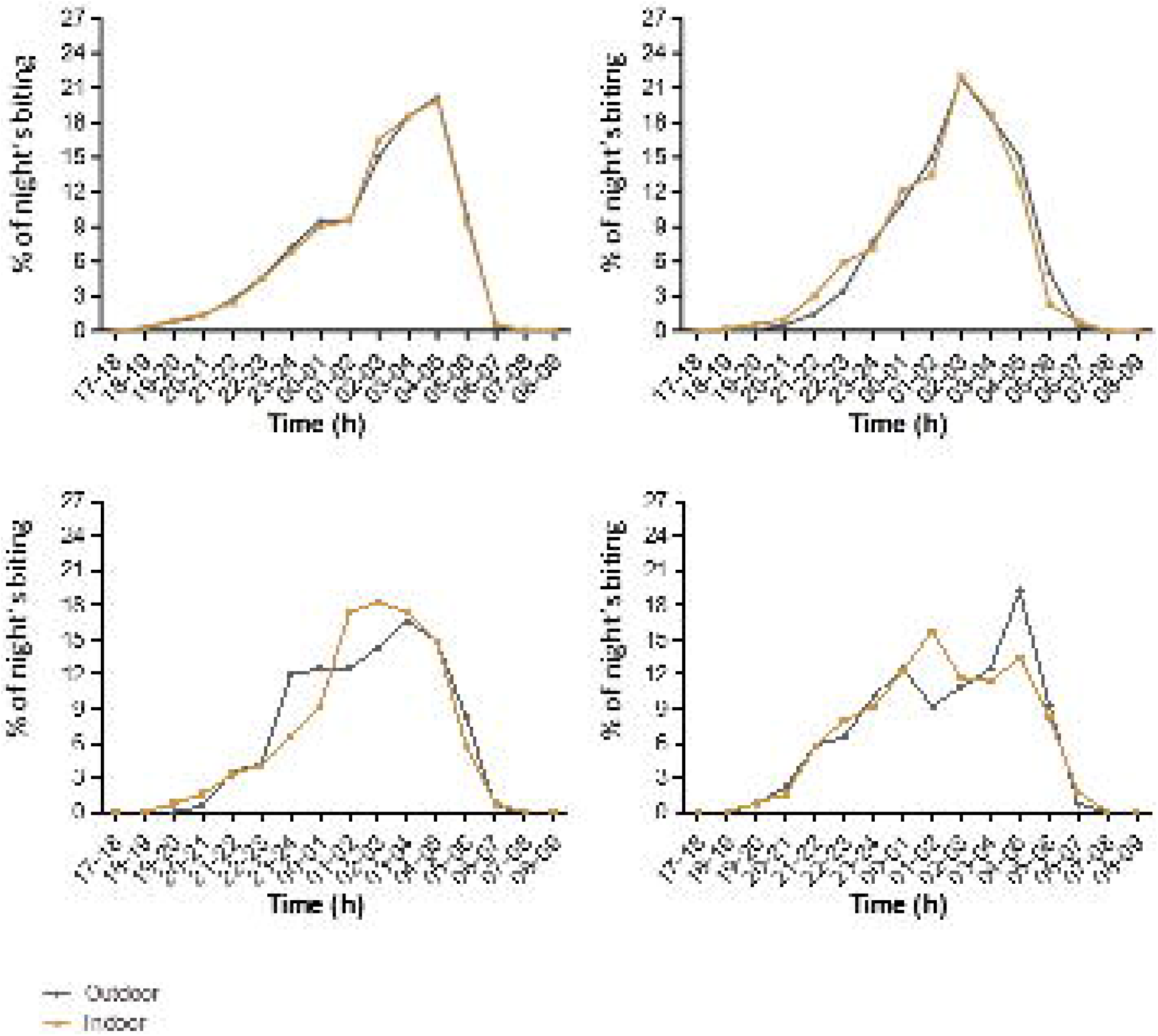

## Infection to *P. falciparum* sporozoite and entomological inoculation rates

*Plasmodium falciparum* sporozoite infection rates per species and surveys are presented in Table 3. The *P. falciparum* sporozoite rate in malaria vector populations was 7.41% (95% IC: 5.47-9.98), 8.98% (95% IC: 5.52-14.29), 4.41% (95% IC: 1.51-12.18) and 1.10% (95% IC: 0.06-5.96) in September-October 2016, November-December 2016, February-March 2017 and April-May 2017, respectively (Table 3). There were no significant differences in the *P. falciparum* sporozoite rates between malaria vector species (χ^2^=7.5, df=3, P=0.057) and surveys (χ^2^=4.9, df=3, P=0.179). The *P. falciparum* sporozoite rates did not differ significantly depending on whether malaria vectors were collected indoor or outdoor (χ^2^=0.0796, df=1, P=0.78).

**Table 3:** *Plasmodium falciparum* sporozoite rates in a subset of *Anopheles* vectors

| Survey | species, group or complex | N positive for Pf | N tested | SR (%) [95%CI] |
| --- | --- | --- | --- | --- |
| September-October 2016 | *Anopheles gambiae s.s.* | 29 | 406 | 7.14 [5.02-10.07] |
|  | *Anopheles coluzzii* | 1 | 10 | 10.00 [0.51-40.42] |
|  | *Funestus* Group | 8 | 47 | 17.02 [8.89-30.14] |
|  | *Anopheles nili* complex | 1 | 63 | 1.59 [0.08-8.46] |
|  | **Total** | **39** | **526** | **7.41 [5.47-9.98]** |
| November-December 2016 | *Anopheles gambiae s.s.* | 14 | 160 | 8.75 [5.28-14.15] |
|  | *Anopheles coluzzii* | 1 | 5 | 20.00 [1.03-62.45] |
|  | *Funestus* Group | 0 | 1 | 0.00 [0.00-94.87] |
|  | *Anopheles nili* complex | 0 | 1 | 0.00 [0.00-94.87] |
|  | **Total** | **15** | **167** | **8.98 [5.52-14.29]** |
| February-March 2017 | *Anopheles gambiae s.s.* | 3 | 42 | 7.14 [2.46-19.01] |
|  | *Anopheles coluzzii* | 0 | 26 | 0.00 [0.00-12.87] |
|  | *Funestus* Group | - | 0 | - |
|  | *Anopheles nili* complex | - | 0 | - |
|  | **Total** | **3** | **68** | **4.41 [1.51-12.18]** |
| April-May 2017 | *Anopheles gambiae s.s.* | 0 | 55 | 0.00 [0.00-6.53] |
|  | *Anopheles coluzzii* | 1 | 36 | 2.78 [0.14-14.17] |
|  | *Funestus* Group | - | 0 | - |
|  | *Anopheles nili* complex | - | 0 | - |
|  | **Total** | **1** | **91** | **1.10 [0.06-5.96]** |

N: number of mosquitoes; SR: Sporozoite rate; Pf: *Plasmodium falciparum*, [95%CI]: 95% Wilson's confidence interval.

Overall, an unprotected individual living in the Korhogo area was estimated to receive an average of 9.04, 0.63, 0.06 and 0.12 infected bites per night in September-October 2016, November-December 2016, February-March 2017 and April-May 2017, respectively. In September-October 2016, an estimated 0.69 infected bites per night (i.e. 20.7 per month) occurred between 17:00 to 22:00. The average EIR recorded in this study was 2.46 infected bites per man per night.

## Frequencies of the *kdr* and *ace-1* alleles

The frequency of the *kdr* allele in *An. gambiae s.s*. population (frequency range: 90.70–100%) was very high throughout the study periods (Table 4). It was significantly higher in September-October 2016 than in February-March 2017 (P < 0.001) and April-May (P < 0.001). The frequency of the *kdr* allele in *An. gambiae s.s*. population in September-October 2016 did not differ from that of November-December 2016 (P = 1.00). The frequency of the *kdr* allele in *An. coluzzii* population ranged from 55.6% to 100% during the four surveys (Table 4). It was significantly higher in September-October 2016 than in February-March 2017 (P < 0.001) and April-May 2017 (P < 0.001). The frequency of the *kdr* allele in *An. coluzzii* population did not vary significantly between September-October 2016 and November-December 2016 (P = 1.00). The *kdr* allele was more frequent in *An. gambiae s.s*. than in *An. coluzzii* population (P < 0.001). We found significant variations of the frequency of the *kdr* allele in *An. gambiae s.s*. (P< 1×10^−4^) and *An. coluzzii* (P=0.020) populations among the 6 villages. For each survey, genotype frequencies of *kdr* in the *An. gambiae s.s* and *An. coluzzii* populations from each of the 6 villages were not different from the Hardy-Weinberg expectations (P>0.05).

**Table 4:** *Kdr* mutation frequencies in a subset of *Anopheles gambiae s.l*.

|  |  | RR | RS | SS | Total | Allelic frequency (%) [95%CI] |
| --- | --- | --- | --- | --- | --- | --- |
| September-October | *Anopheles gambiae* s.s. | 402 | 1 | 0 | 403 | 99.88 [99.30-99.99] |
|  | *Anopheles coluzzii* | 9 | 1 | 0 | 10 | 95 [76.39-99.74] |
| November-December | *Anopheles gambiae* s.s. | 160 | 0 | 0 | 160 | 100 [98.81-100] |
|  | *Anopheles coluzzi* | 5 | 0 | 0 | 5 | 100 [72.25-100] |
| February-March | *Anopheles gambiae* s.s. | 37 | 4 | 2 | 43 | 90.70 [82.70-95.21] |
|  | *Anopheles coluzzii* | 11 | 11 | 4 | 26 | 63.46 [49.87-75.20] |
| April-May | *Anopheles gambiae* s.s. | 52 | 2 | 2 | 56 | 94.64 [88.80-97.52] |
|  | *Anopheles coluzzii* | 12 | 16 | 8 | 36 | 55.56 [44.09-66.46] |

RR: homozygous resistant, RS: heterozygote, SS: homozygous susceptible, [95%CI]: 95% Wilson's confidence interval.

The *ace-1* frequency in *An. gambiae s.s*. population ranged from 15.18 to 30.05 % throughout the study periods (Table 5). The frequency of the *ace-1* allele in *An. gambiae s.s*. population was significantly higher in September-October 2016 than in November-December 2016 (P= 0.021), February-March 2017 (P= 0.026) and April-May 2017 (P<0.001). The frequency of the *ace-1* allele in *An. coluzzii* population (frequency range: 15.18–30.05 %) did not vary significantly among the four surveys (P=0.278). The *ace-1* allele was significantly more frequent in *An. gambiae s.s*. than in *An. coluzzii* population (P < 1×10^−4^). The frequency of the *ace-1* allele in *An. gambiae s.s*. populations varied significantly among the 6 villages (P< 1×10^−4^). The frequency of the *ace-1* allele in *An. coluzzii* population did not vary significantly among the 6 villages (P=0.211). For each survey, genotype frequencies of *ace-1* in *An. gambiae* and *An. coluzzii* populations from each of the 6 villages did not differ from the Hardy-Weinberg proportions (P>0.05).

**Table 5:** *Ace1* mutation frequencies in a subset of *Anopheles gambiae s.l*.

|  |  | RR | RS | SS | Total | Allelic frequency (%) [95%CI] |
| --- | --- | --- | --- | --- | --- | --- |
| September-October | *Anopheles gambiae s.s.* | 41 | 156 | 199 | 396 | 30.05 [26.96-33.33] |
|  | *Anopheles coluzzii* | 0 | 1 | 9 | 10 | 5 [0.26-23.61] |
| November-December | *Anopheles gambiae s.s.* | 15 | 43 | 101 | 159 | 22.96 [18.67-27.88] |
|  | *Anopheles coluzzii* | 0 | 0 | 5 | 5 | 0 [0-27.75] |
| February-March | *Anopheles gambiae s.s.* | 2 | 12 | 29 | 43 | 18.60 [11.79-28.10] |
|  | *Anopheles coluzzii* | 0 | 0 | 26 | 26 | 0 [0-6.88] |
| April-May | *Anopheles gambiae s.s.* | 3 | 11 | 42 | 56 | 15.18 [9.70-22.97] |
|  | *Anopheles coluzzii* | 0 | 1 | 35 | 36 | 1.39 [0.07-7.46] |

RR: homozygous resistant, RS: heterozygote, SS: homozygous susceptible, [95%CI]: 95% Wilson's confidence interval.

## Discussion

*Anopheles gambiae s.l*. is by far the main malaria vector in the Korhogo area, irrespective of the season, but its abundance varied significantly according to the season. The risk of being bitten by malaria vector mosquitoes was found to be up to 200-fold higher in September-October 2016 (rainy season) than in February-March 2017 (dry season). These results are consistent with the bionomics of *An. gambiae s.s*. Indeed, the breeding habitats of this species are known to increase in number and productivity during the rainy season but almost disappear during the dry season [31]. In September-October 2016 (rainy season), an unprotected individual in the Korhogo area received an estimated ~9 infected bites per night (i.e. 270 per month). Korhogo is a rural area where several cultures including rice are cultivated. Rice paddies are strongly associated with very high densities of malaria vectors [20]. In this study, we found a mean annual EIR of 897.9 infected bites per person per year. Research conducted so far in Africa have reported mean annual EIRs ranging from < 1 to > 1,000 infected bites per person per year [3]. Therefore, Korhogo ranks at the top level among malaria endemic areas in terms of the intensity of transmission.

Our results revealed that malaria vector populations were significantly exophagic in both the rainy and dry seasons, with no significant difference in outdoor biting rates between seasons. Furthermore, we found that indoor and outdoor biting activities of *An. gambiae s.l*. peaked later (4-5 am) in September-October 2016 than in the three other surveys. The influence of genetic factors in the biting behaviors of malaria vectors remains poorly understood [27]. However, some studies have shown that *Anopheles* vectors can adjust their biting behavior in response to a small change in environmental parameters such as temperature [9,32]. During the 4 surveys, we recorded environmental data (temperature, humidity, atmospheric pressure, light intensity) in all the sites during the collections. These data would help establish a possible link between environmental variables and the biting behaviors of vectors. It is more and more obvious that LLINs alone are unable to eliminate malaria in high malaria burden countries [19]. In this study, we found that an estimated 0.69 infected bites (i.e. 20.7 per month) occurred daily in September-October 2016 between 17:00 to 22:00 when human populations are potentially unprotected by ITNs. Nonetheless, our results need to be adjusted to the behaviors of the local human populations in terms of sleeping hours, outdoor activities and ITNs use in order to more accurately measure human exposure to *Plasmodium* sp.-infected mosquito biting in the Korhogo area [29]. We will address this research question in another publication using additional data collected in the frame of the REACT project.

Consistent with recent studies carried out in the Korhogo area [6,25], we found very high frequencies of the *kdr* mutation in the *An. gambiae* complex population during the four surveys. The *kdr* mutation has almost reached fixation in *An. gambiae s.s*. population across Côte d’Ivoire [6]. We identified the *ace-1* allele in the *An. gambiae* complex populations at frequencies ranging from 0 to 30.05% during the four surveys. A recent study has shown that *An. gambiae s. l* mosquitoes from Korhogo are highly resistant to pyrethroids, organochlorides, and carbamates but susceptible to organophosphates [6]. Accordingly, the use of organophosphates may represent a valuable strategy for the control of this high insecticide resistant vector population. However, this approach would select genotypes bearing *ace-1* mutation which display resistance to organophosphates although the *ace-1* mutation confers a lower resistance level to organophosphates than carbamates [11]. In our study, we found spatial and temporal variations of the *kdr* and *ace-1* frequencies in the *An. gambiae* complex population. The frequencies of insecticide resistance alleles in malaria vector populations are known to be influenced by the selective pressures by the use of insecticides in agriculture and public health as well as by the fitness costs associated with resistance alleles in absence of insecticide [1,10,23,28,41]. Both the *kdr* and *ace-1* alleles were significantly more frequent in *An. gambiae s.s*. than in *An. coluzzii* population. These differences in *kdr* and *ace-1* frequencies between *An. gambiae s.s*. and *An. coluzzii* have been observed in many areas of Africa [7,8,12].

This study demonstrated that malaria vectors were more likely to feed outdoors than indoors and the frequencies of the *kdr* allele were very high in *An. gambiae s.l*. populations. Moreover, we found that the risk of malaria transmission was significant when people are potentially active. Therefore, malaria transmission would still continue in the Korhogo area even with full coverage of LLINs. A large-scale multi-country study published recently showed conclusively that LLINs are still protective against malaria despite widespread resistance to pyrethroid [22]. However, malaria control in high endemic areas such as Korhogo needs to be strengthened with complementary tools to alleviate the burden of the disease. Therefore, efforts need to be dedicated to the evaluation of the effectiveness of promising tools to be used in combination with LLINs in high endemic areas. The REACT’s project would provide evidence for the impact of four potential strategies on clinical malaria and transmission in two high endemic areas of Burkina-Faso and Côte d’Ivoire. Two of these strategies (larviciding with *Bti* and use of Ivermectin in cattle and other small ruminants) are at late-stage of development and the two others (intensive communication for human behavioral changes and indoor residual sprayings of insecticide) are already available in the vector control arsenal against malaria.

## Conclusions

Malaria transmission in the Korhogo area was mainly due to *An. gambiae s.s*. and *An. coluzzii* while *An. funestus* group species and *An. nili* complex species played minor roles. The frequencies of the *kdr* mutation in the *An. gambiae* complex population were very high whereas the *ace-1* frequencies were moderate. Malaria vectors in Korhogo were more exophagic. The intensity of malaria transmission is extremely high in the Korhogo area, especially during the rainy season. Malaria control in such highly endemic areas needs to be strengthened with complementary tools in order to reduce the burden of the disease.

## Acknowledgements

We thank all the mosquito collectors and supervisors for commitment in the field. We also thank Koné Aboubacar, Azaibou Akoliba Patrice, Dosso Youssouf, Koffi Serges, Konan Koffi Guillaume and Wolie Rosine for their technical assistance.

